# Gaps in the global protection of terrestrial genetic diversity

**DOI:** 10.1101/2024.08.23.609342

**Authors:** Jana Tabea Schultz, Jonas Geldmann, Spyros Theodoridis, David Nogues-Bravo

## Abstract

In recent decades, increased anthropogenic impact has led to a global decline in genetic diversity. Before the Kunming-Montreal Global Biodiversity Framework (2022), the absence of international consensus on how to directly assess and monitor genetic diversity, hampered large-scale conservation efforts. Scarcity of assessable genetic data has hindered the evaluation of conservation policies in safeguarding genetic diversity. This study presents the first global approach for evaluating the protection of genetic diversity. By examining the global distribution of mammalian intraspecific mitochondrial DNA and protected area coverage, we identify regions with high genetic diversity and insufficient protection coverage, e.g. regions of critical importance for biodiversity in the Brazilian Atlantic Forest. Additionally, we estimate the impact of global change scenarios on genetically diverse regions with a low degree of protection, revealing high vulnerability of areas in Central Africa. Nonetheless, integrating robust analysis into conservation planning remains challenging. Incorporating Macrogenetics into conservation planning holds the potential to reverse biodiversity decline.

## 1 Introduction

Human impact on the biosphere has accelerated over the last decades, significantly altering over 75% of the Earth’s terrestrial surface (IPBES, 2019). This has led to an increased risk of extinction; around 25% of assessed species are being classified as endangered. As the anthropogenic pressure on the environment continues to increase, the loss of biological diversity will further accelerate (IPBES, 2019), marking the path toward a possible sixth mass extinction. This biodiversity decline threatens ecosystem functions and compromises the integrity of the biosphere, ultimately jeopardizing the well-being of human societies (Richardson et al., 2023).

Biodiversity is defined as the variety of life on Earth and encompasses genetic, species, and ecosystem diversity as three inter-connected elements. However, genetic diversity has often been overlooked in international policy agreements (Hoban et al., 2021a; Laikre et al., 2020). In an effort to enhance the conservation of genetic diversity, the Convention on Biological Diversity’s (CBD) recently agreed on the Kunming-Montreal Global Biodiversity Framework (GBF), which calls for the preservation and restoration of genetic diversity within and between populations of native, wild and domesticated species (i.e. Target 4; CBD COP, 2022), but the lack of available data challenges the operationalization of these aspirations. This is for example illustrated by GBF’s monitoring framework, suggesting that measures of effective population size could be used as a crude proxy of genetic diversity as measuring actual genetic diversity remain difficult, costly, and impractical (Hoban et al., 2021b) with no scientific agreement on how to directly assess or monitor genetic diversity (Forcina & Leonard, 2020; Hoban et al., 2021a).

Consequently, biological diversity has been mainly inferred as an aspect of species richness, abundance metrics, and habitat variation, leaving genetic diversity insufficiently monitored and aggregated for large-scale analysis and conservation approaches (Hoban et al., 2022). However, recent technology advances, improved data infrastructures, and open assess initiatives have paved the way for an increase in genetic studies at large spatial scale (McCallen et al., 2019). These advancements have been critical for the development of the new discipline “Macrogenetics”, enabling the exploration of the conservation of genetic diversity at larger spatial scales (Hoban et al., 2021b; Leigh et al., 2021). The emerging global repositories of geo-referenced genetic samples have enabled studies of the global geography of intraspecific genetic diversity (Leigh et al., 2021; Miraldo et al., 2016; Theodoridis et al., 2020) revealing that the decline of genetic diversity has likely already surpassed 10% (Exposito-Alonso et al., 2022). This is concerning as intraspecific genetic diversity, or the variation in a DNA sequence among individuals within a species, is essential for species’ adaptation to changing environmental conditions (Nonić & Šijačić-Nikolić, 2021; Theodoridis et al., 2020).

Protected areas have long been regarded as one of the most important tools for safeguarding biodiversity, with the Global protected area estate currently protecting 16.02% of Earths terrestrial surface (UNEP-WCMC & IUCN, 2023). While this is an impressive achievement, the GBF raises the ambitions, with Target 3 stating that 30% of the land and the oceans will be protected by 2030 (CBD COP, 2022). This provides an important impetus for ensuring that future expansions effectively target the area of most importance for biodiversity. The effectiveness of protected areas in terms of covering the most important places for biodiversity have traditionally been assessed using species richness, endemism, and vulnerability (Daru et al., 2019; Margules & Pressey, 2000). However, focussing solely on species-level metrics may overlook other vital aspects of biodiversity, such as phylogenetic diversity (Aguilar-Tomasini et al., 2021; Daru et al., 2019) and the diversity of ecosystems (Keith et al., 2015). Additionally, the absence of molecular data in the global conservation planning process means that we lack crucial information on the adaptive potential of species (Andrello et al., 2022; Nielsen et al., 2022). This shortfall may compromise the long-term success of protected areas in preserving biodiversity (Dudley, 2008; Rodrigues & Cazalis, 2020). It is imperative, therefore, to integrate recent findings of global patterns of genetic diversity into identifying biodiversity importance areas and applying Systematic Conservation Planning at all scales.

Here, we illustrate how the integration of Macrogenetics into area-based conservation provides a pathway to evaluate the global network of protected areas’ ability to safeguard intraspecific genetic diversity. Specifically, we assess the extent to which the global network of protected areas, excluding Other Effective Area-based Conservation Measures (OECMs), geographically covers the mitochondrial genetic diversity of terrestrial mammals. To achieve this, we intersect terrestrial protected areas (UNEP-WCMC & IUCN, 2021) and the geographical distribution of intraspecific genetic diversity of mammal species – the only taxonomic group for which sufficient amount of Macrogenetic data is available at the global scale (Theodoridis et al., 2020): this highlights the unique and complimentary information genetic diversity provides for future conservation planning. Our aim is to evaluate the representation of intraspecific genetic diversity within protected areas globally and identify regions of the world that are in urgent need of protection due to predicted high exposure to environmental change under alternative Shared Socioeconomic Pathways (SSPs).

## 2 Methods

### 2.1 Data on genetic diversity

Information about the geographical distribution of genetic diversity was sourced from Theodoridis et al. (2020).Briefly, Theodoridis et al. (2020) modelled globally genetic diversity using the two most extensively used genetic markers, cytochrome oxidase I (*CO1*) and cytochrome b (*cytb*). Modeled biodiversity data is essential in biogeography for providing comprehensive global coverage, helping to reduce geographical bias inherent in observed data, is cost-effective and it supports informed decision-making in conservation and policy, filling crucial gaps due to the Wallacean shortfall (Hortal et al. 2015).

Georeferenced genetic sequences were obtained from GeneBank and BOLD in May 2017, using the API tool provided by GeoNames (http://api.geonames.org). The generated genetic database contained 46,965 mitochondrial sequences: 24,395 sequences for *cytb* and 22,570 sequences for *CO1*. Further, the sequence data was aligned, and sequences were excluded if their geographical coordinates did not coincide with the known native ranges of species defined by the IUCN database. To map the distribution of genetic diversity, the globe was divided into an equal-area grid (Behrmann cylindrical equal-area projection) with a cell size of 385.9km × 385.9km representing 148,953km^2^ area grid-cells (see Theodoridis et al. 2020 for methodological protocols).

Theodoridis et al. (2020) predicted the global distribution of genetic diversity using phylogenetic diversity and temperature trend as explanatory variables for *CO1* and solely phylogenetic diversity for *cytb* using linear models (see Theodoridis et al. 2020 for details). Genetic Diversity (GD) was measured using nucleotide diversity which is, the average number of nucleotide differences per site in a pairwise sequence comparison across species within each grid-cell (Miraldo et al., 2016).

### 2.2 Data on protected areas

Data on the global spatial distribution of protected areas were obtained from the World Database on Protected Areas (WDPA; December 2021 released), which is the primary reference source of spatial data for protected areas worldwide and the official data-repository for reporting towards the CBD. Only protected areas reported as polygons and defined as terrestrial were selected, leaving us with 252,431 protected areas. We also excluded OECMs for which globally consistent data does not exist. We then calculated the percentage coverage of terrestrial land by protected areas within each grid cell using a grid with the same spatial resolution and projection as our genetic maps.

### 2.3 Protection of genetic diversity

To counteract the bias towards the equator due to the global distribution of biodiversity, each grid cell was assigned to one of the 14 bioclimatic regions based on the most represented biome in the cell. The biomes were defined by Dinerstein et al. (2017).

We examined the relationship between genetic diversity and coverage by protected areas using linear regression model (LM) at the global (i.e. across grid-cells) and at the biome level, whereat genetic diversity was normalized using square root transformation.

Further, we compared the conservation efforts between the bioclimatic regions by calculating the mean and standard errors of protection coverage and genetic diversity for each region.

### 2.4 Discrepancy between genetic diversity and species richness

Under the assumption that species richness correlates with genetic variety, protecting species richness may also safeguard genetic diversity. To assess the discrepancy between genetic diversity and species richness, we first subset the GD dataset by keeping only data-rich grid cells (cells with a minimum of 55 sequences for *cytb* and 278 sequences for *CO1*; for more details see methods in Theodoridis et al. (2020)) to overcome the differences in data availability (number of species used for estimating GD compared to the total number of species naturally occurring in each grid cell estimated from IUCN range maps). We transformed GD to normality using its square root before running a linear model using species richness as independent variable for both genetic markers, as well as temperature trend for *CO1*, and observed genetic diversity as response. We then mapped the residuals of the regression to identify regions with higher or lower genetic diversity as predicted by species richness.

### 2.5 Exposure of unprotected genetic diversity to land-use change

To examine the exposure of unprotected genetically diverse areas under future environmental changes, we combined information on the spatial correlation of genetic diversity and conservation coverage with the prediction about future agricultural expansion under the four main SSP scenarios: SSP126, 245, 370, and 585. Data on land-use change under the alternative SSP scenarios were sourced from the Land-Use Harmonization (LUH2) project (https://luh.umd.edu/data.shtml). To this end, we selected sites representing the 20% most genetically diverse areas but show less than 10% coverage by protected areas.

## 3 Results

### 3.1 Protection of genetic diversity

Our LM did not show any correlation between the percentage coverage by protected areas and genetic diversity at the global level for any of the two genetic markers (*cytb*: R^2^ = 0.0006, *p* = 0.37, 95% CI [-80.95, 30.04]; *CO1*: R^2^ = 0.0009, *p* = 0.26, 95% CI [-70.1, 19.21]). Similarly, we did not observe any significant correlation between the two genetic markers within bioclimatic regions (Table S1). However, our findings indicate that while several areas with high genetic diversity, such as the Amazon, Central America, and Eastern Africa have extensive protected area coverage, the current global network of protected areas has significant gaps, leaving numerous genetically diverse areas unprotected (Figure 1). This is particularly the case for areas of North America outside of the west coast and Alaska, Central and East Asia, as well as regions of critical importance for biodiversity in the Brazilian Atlantic Forest, Southern Africa, and South-East Asia (Figure 1). At the bioclimatic region level, we show that biomes hosting the highest genetic diversity, including tropical, and subtropical grasslands, savannahs, and shrub lands or tropical and subtropical moist broadleaf forests, are not adequately protected (Figure S3).

**Figure 1:**
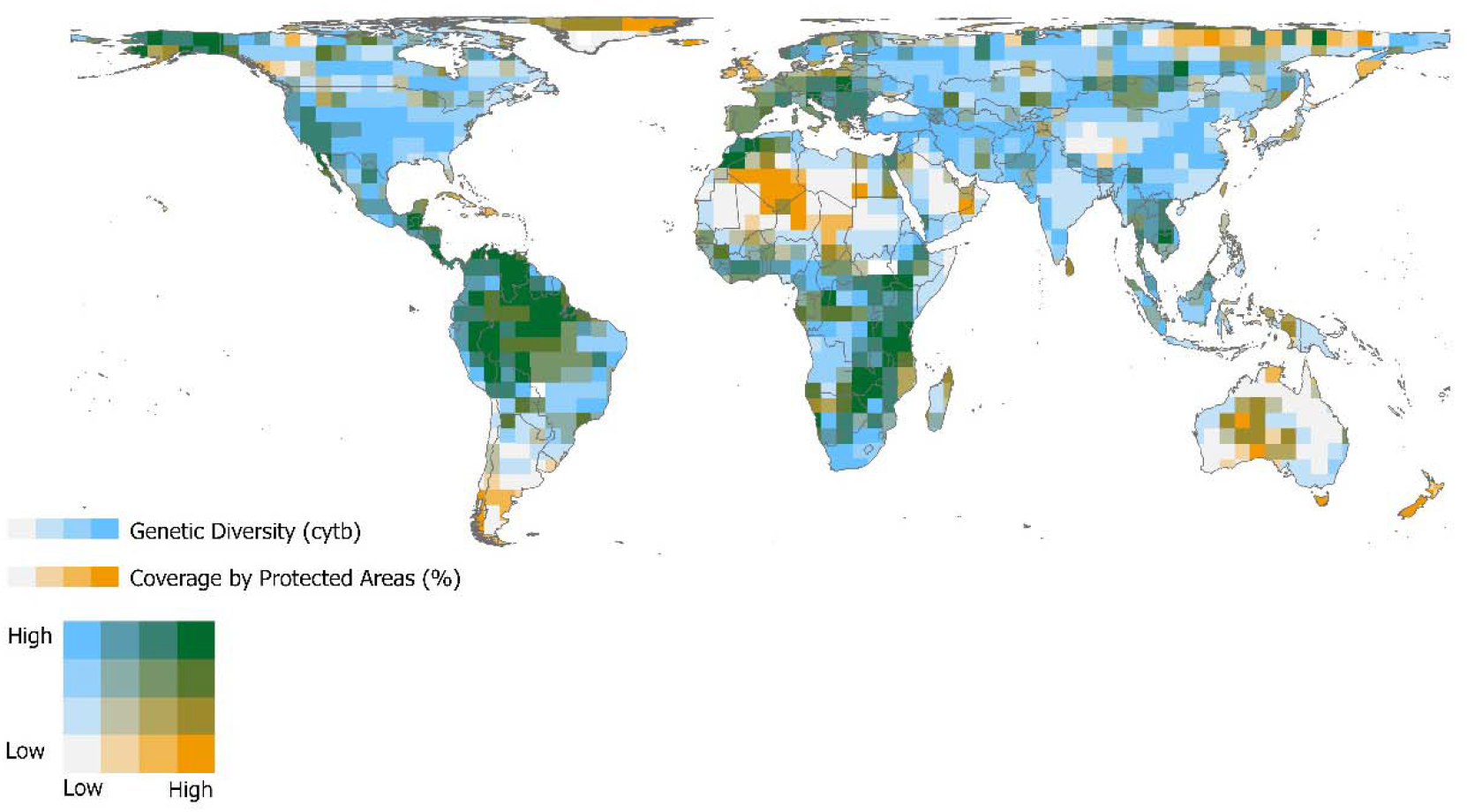
Regions of the planet hosting high genetic diversity but lacking coverage by protected areas. Coverage of predicted genetic diversity (*cytb*) by the global network of protected areas based on the bioclimatic regions (Figure S2). Each cell is representing 148,953 km^2^ area, as this cell size is most suitable for the calculation of genetic diversity using the formular by Tajima (1993). Every grid cell was assigned to the most represented bioclimatic region in the cell. The coverage by protected areas is represented in percentage and was grouped based on conservation policy targets (< 10%, 10 – 17%, 17 – 30%, > 30%). Hence, white colour represents a coverage by protected areas lower than 10 %, orange shows a coverage over 30%. Genetic diversity data was categorized by quantiles. Map showing the genetic diversity based on *CO1* can be found in the supplementary material (Figure S1).

### 3.2 Discrepancy between genetic diversity and species richness

Our results indicate that genetic diversity and species richness overall are correlated (*cytb*: R^2^ = 0.34, *p* = 2e-16, 95% CI [0.0002, 0.0003] ; *CO1*: R^2^ = 0.3, *p* = 0.03, 95% CI [1.13, 0.0002]). However, our findings also reveal regions where genetic diversity is higher than species richness indicates, e.g., Southern Cone and South Africa, and where species richness is not a suitable indicator for genetic diversity (Figure 2; Figure S4).

**Figure 2:**
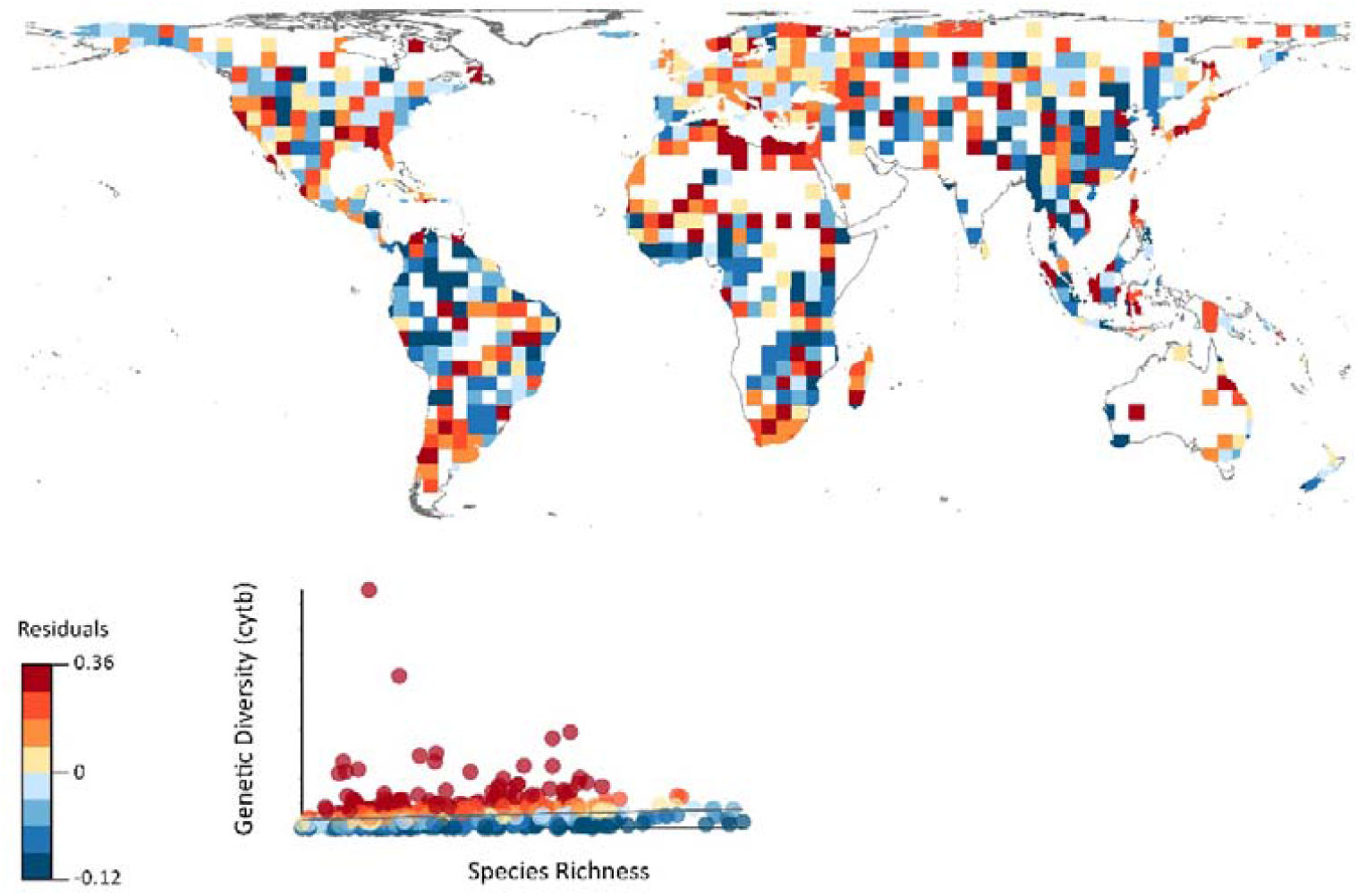
The geographical distribution of the residuals between observed genetic diversity (*cytb*; 719 cells) and species richness. Dark blue shows areas where genetic diversity is lower than predicted by species richness (i.e. negative residuals). Dark red shows regions where genetic diversity is higher than predicted by species richness (i.e. positive residuals). Map showing the geographical distribution of the residuals between observed genetic diversity (*CO1*; 380 cells) and species richness can be found in the supplementary material (Figure S4).

### 3.3 Future global change impacts in unprotected areas

Our results reveal that numerous highly genetically diverse areas are inadequately protected (with less than 10% coverage). Under the SSP3 scenario, also known as regional rivalry scenario, genetically diverse areas will overall experience the highest pressure of agriculture expansion. Especially regions in Africa will be highly affected by predicted agriculture expansion of up to 43.1% (Figure 3, Figure S4). The SSP5 scenario, fossil-fuelled development, also poses a high threat to genetically rich regions, especially in Central Africa with a predicted agriculture expansion up to 47.9%. However, on the global scale the majority the sites will experience only experience a minor expansion of agricultural land (< 1.6%) or even a reduction (> -10%).

**Figure 3:**
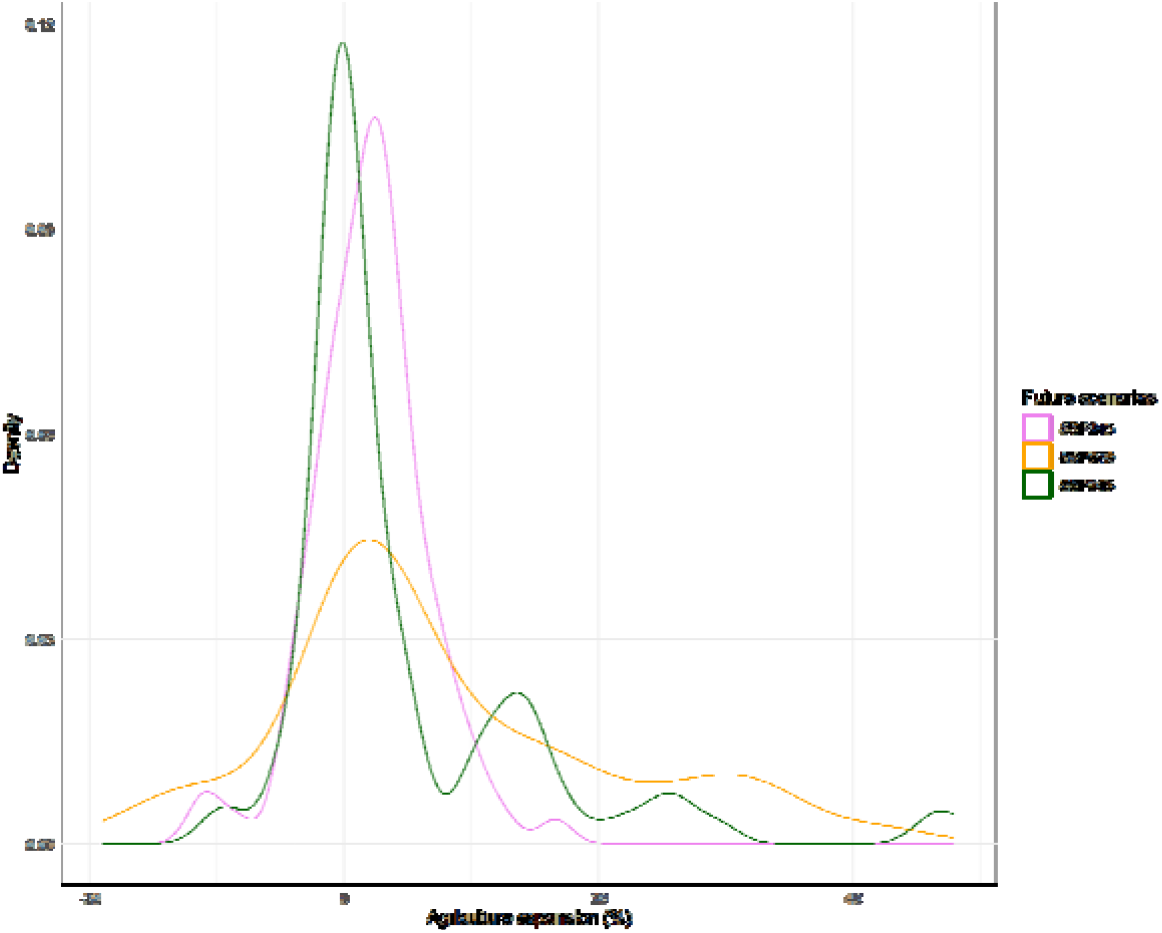
Exposure of genetically diverse areas to agricultural expansion under different SSP scenarios. We assess sites holding 20% of the most diverse populations, but with low protection coverage (< 10%). We do not consider SSP126 as no changes in the size of agricultural land are predicted under this scenario (Figure S5).

## 4 Discussion

Our results here provide the first estimation of the degree of protection of mammal intraspecific genetic diversity globally and underscore the urgency of integrating genetic diversity into SCP. Through this, we show that while some areas of importance for genetic diversity are among the best protected parts of the world, many of the most genetically diverse areas are lacking protection. Importantly, we also show that patterns of genetic diversity are not capture but patterns of species richness and therefore protecting species richness may not necessary protect genetic diversity. Thus, in line with previous studies (Daru et al., 2019; Mazel et al., 2018), our research demonstrates that relying solely on species-level diversity metrics cannot reliably function as a surrogate for other features of biodiversity. Recent studies analysed the potential of the integration of genetic data in SCP and identified possible frameworks for the implementation of these data in conservation (Andrello et al., 2022; Nielsen et al., 2022; van Oppen & Coleman, 2022). However, attempts to integrate genetic data on a global scale are still hindered by the lack of available and accessible DNA data.

Although mtDNA may be so far the only type of genetic data rich enough for a global view of the distribution of genetic diversity for thousands of species, as it is the most frequently used genetic marker for species identification and biodiversity monitoring (Krishnamurthy et al., 2012), the use of mtDNA in connection with biodiversity conservations assessment should be interpreted with care. Genetic diversity data based on mitochondrial markers is difficult to compare because mtDNA mutation rates highly vary across taxa (Schmidt & Garroway, 2021). Nevertheless, owing to technological advancement, there is an ever-increasing availability of genomic data across space and taxa (Leigh et al., 2021). Genomic data for thousands of species and populations across the planet will provide new opportunities to improve our understanding of the relation between different biodiversity metrics and the spatial distribution of genetic variability (Schmidt & Garroway, 2021; Theodoridis et al., 2020), and also to enhance the feasibility of the integration of genetic data into conservation planning. Incorporating large genetic datasets into conservation approaches, such as SCP, can help to identify areas that are critical for conservation (Leigh et al., 2021).

Developing robust analysis incorporated in SCP is still challenging: here we identify and discuss three challenges ahead: the genomic Wallacean shortfall, the baseline shortfall, and the metric shortfall. Recently, those challenges have received significant attention and different pathways have been proposed to tackle them (Andrello et al., 2022; Hoban et al., 2022; Miraldo et al., 2016). Nevertheless, we are still far from having the information and metrics that we already have for other facets of biodiversity (e.g. species distribution, traits, and phylogenies).

The “Wallacean shortfall” refers to the limited knowledge about the geographical distribution of species (Hortal et al., 2015). In the context of genetic diversity, this shortfall may cause that the distribution of genetic diversity reflects the survey effort rather than actual diversity patterns. Currently, due to globally unequal sample efforts, both for mtDNA and nuclear DNA (Leigh et al., 2021; Theodoridis et al., 2020) there are strong geographical and taxonomical biases in data availability. In particular, highly diverse regions such as subtropical and tropical biomes show substantially lower data availability and sampling efforts (Miraldo et al., 2016; Theodoridis et al., 2020). To address this shortfall, increased effort in data collection and georeferencing are needed, specifically in data-poor regions, such as South America and Central Africa (Leigh et al., 2021). Moreover, the development of models predicting different aspects of genetic diversity in unsampled regions, as the modelled data used for our study (Theodoridis et al., 2020), and the use of contemporary specimens already collected and curated in natural history collections will help to provide a more accurate representation of global genetic diversity pattern.

The “baseline shortfall” refers to the lack of knowledge about past genetic diversity levels for most of the species, impeding the estimation of genetic diversity trends (i.e., declines) over time. Ancient DNA have been proven useful tools to predict past levels of genetic diversity (e.g., for the Late Pleistocene), serving as a baseline for pre-Anthropocene time periods (Jensen et al., 2022; Leigh et al., 2021). However, this approach is limited to species and regions with high fossilization potential. Whole-genome resolution data from contemporary specimens can provide additional information, as they allow, albeit with certain limitations, the estimation of population size over time even in the absence of a DNA (Iannucci et al., 2021; Sato et al., 2020). Utilizing such data can help address the baseline shortfall and enhance our understanding of genetic diversity patterns across various time scales.

The “metric shortfall” defines the absence of an international agreement on what is the most feasible and appropriate genetic diversity metrics for conservation purposes. This topic is the subject of significant debate and its magnitude is beyond the scope of our study (Hoban et al., 2022a; Hoban et al., 2021b; Leigh et al., 2021). However, the implementation of other biodiversity metrices, such as species richness or endemicity level, in spatial conservation approaches have showed that genetic diversity metrics should be able to be estimated for a significant amount of the species present in a specific location, with low-economic cost and comparable across different realms (i.e., marine versus terrestrial). This would enable more effective and efficient conservation strategies and help address the metric shortfall in biodiversity conservation efforts. Our study measures genetic diversity through the analysis of nucleotide diversity. This approach, while not encompassing all aspects of genetic diversity, offers a valuable indicator of intraspecific genetic variation, as demonstrated in Theodoridis et al. (2020). We acknowledge that our modelled does not directly measure the abundance component of genetic diversity, but it provides a robust proxy for it. As genomic technologies advance, our approach can be refined to incorporate more nuanced measurements, such as genetic uniqueness and allele abundance metrics. Nevertheless, our current methodology is a critical step in understanding and conserving the genetic facet of biodiversity, especially given the Wallacean shortfall in genetic data highlighted by Theodoridis et al 2020.

By overcoming these shortfalls, genetic data could become increasingly relevant and applicable for SCP, playing a crucial role in identifying evidence-based global conservation priorities (Zizka et al., 2022). Including conservation prioritization in Marcogenetic studies has the potential to enhance the effectiveness of conservation efforts as conservation priority areas identified using intraspecific genetic data from multiple species provide more effective conservation solutions than areas identified for single species or on the basis of traditional taxonomic criteria (Paz-Vinas et al., 2018). Thus, we advocate for genetic diversity to be given equal importance in spatial conservation strategies, on a par with other crucial factors such as species diversity. By doing so, we can better maintain biodiversity, protect the integrity of the biosphere, and ensure the well-being of human society.

Only by integrating Macrogenetics in conservation planning there is a chance to reverse the current biodiversity decline as genetic diversity gives species the potential to persist under environmental change as expected in the face of anthropogenic action (Andrello et al., 2022). Thus, progress towards increased standardization in measuring and monitoring genetic diversity is necessary if we are to deliver on the Kunming-Montreal Global Biodiversity Framework’s ambitions to fully integrate genetic diversity in assessing the true health of our planet.

## Acknowledgment

JS acknowledge financial support from the project ‘OR ELSE: Operational Recommendations for Ecosystem-based Large-scale Sand Extraction’ with project number NWA.1389.20.097 of the NWA research program ‘Research along Routes by Consortia (ORC)’, which is funded by the Dutch Research Council (NWO).

DNB thanks DFF Independent Research Fund Denmark (grant 8021-00282B: DEMOCHANGE).

## Data Availability

All data supporting the results of this study are already publicly available. The identifiers for all genetic sequences used in this study and the processed data to recreate the figures are available in https://github.com/spyrostheodoridis/Genetic-geography-of-terrestrial-mammals. An interactive exploration of the reported findings is available through the web application https://geneticgeography.com. The raw genetic sequences are available in GeneBank (www.ncbi.nlm.nih.gov/genbank) and BOLD (www.boldsystems.org). Dated phylogenies are available in Dryad (https://datadryad.org/stash/dataset/doi:10.5061/dryad.bp26v20). Data for bioclimatic regions are available in https://ecoregions.appspot.com/. The protected areas database can be found here: https://www.protectedplanet.net/en/thematic-areas/wdpa?tab=WDPA; the data on land-use change under the SSP scenarios were sourced from the Land-Use Harmonization (LUH2) project (https://luh.umd.edu/data.shtml). Maps will be additionally accessible with publication through GitHub before publication.

## Supporting Information

Additional supporting information will be found in the online version of the article at the publisher’s website.

## Code availability

All code for data cleaning and analysis associated with the current version will be made available on GitHub before publication.

