## Supplementary Information for "Gaps in the global protection of terrestrial genetic diversity"

### Figure S1: Protection of genetic diversity by protected areas based on bioclimatic regions

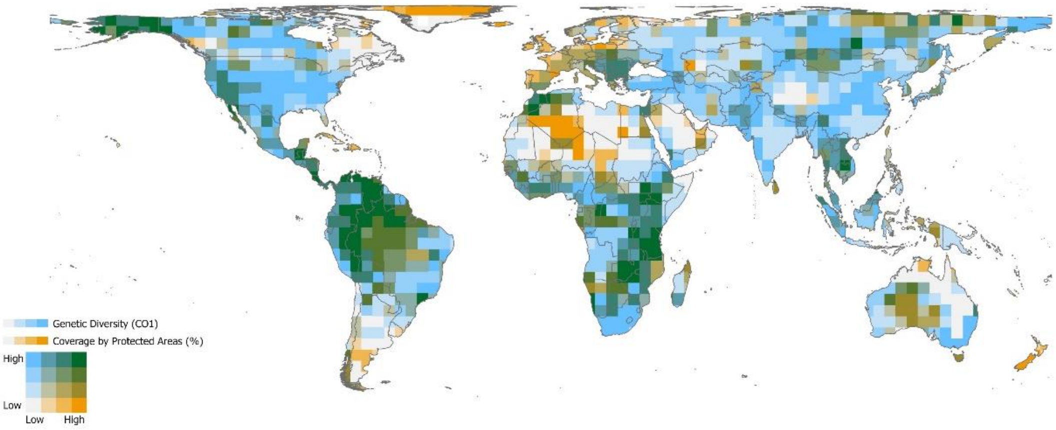


**Figure S1:** Coverage of genetic diversity (CO1) by the global network of protected areas based on the bioclimatic regions. Each cell is representing 148,953 $km^2$ area, as this cell size is most suitable for the calculation of genetic diversity using the formular by Tajima (1993). Every grid cell was assigned to the most represented bioclimatic region in the cell. The coverage by protected areas is represented in percentage and was grouped based on conservation policy targets (< 10 %, 10 – 17 %, 17 – 30 %, > 30 %). Hence, white colour represents a coverage by protected areas lower than 10 %, orange shows a coverage over 30 %. Genetic diversity data was categorized by quantiles.

**Figure S2: The allocation of io bioclimatic regions**

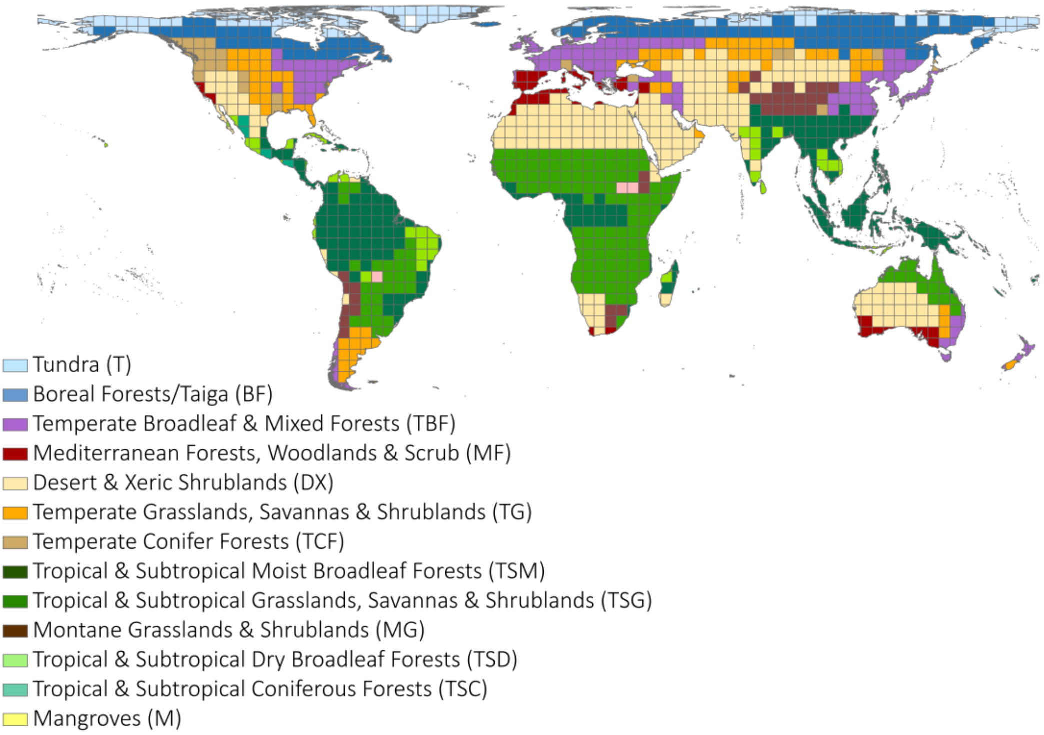


**Figure S2:** Each grid cell was assigned to one bioclimatic regions based on the most represented biome in the cell. The biomes followed the definition of Dinerstein et al. (2017).

**Figure S3: The average coverage by protected areas of genetic diversity in each bioclimatic region**

(a)

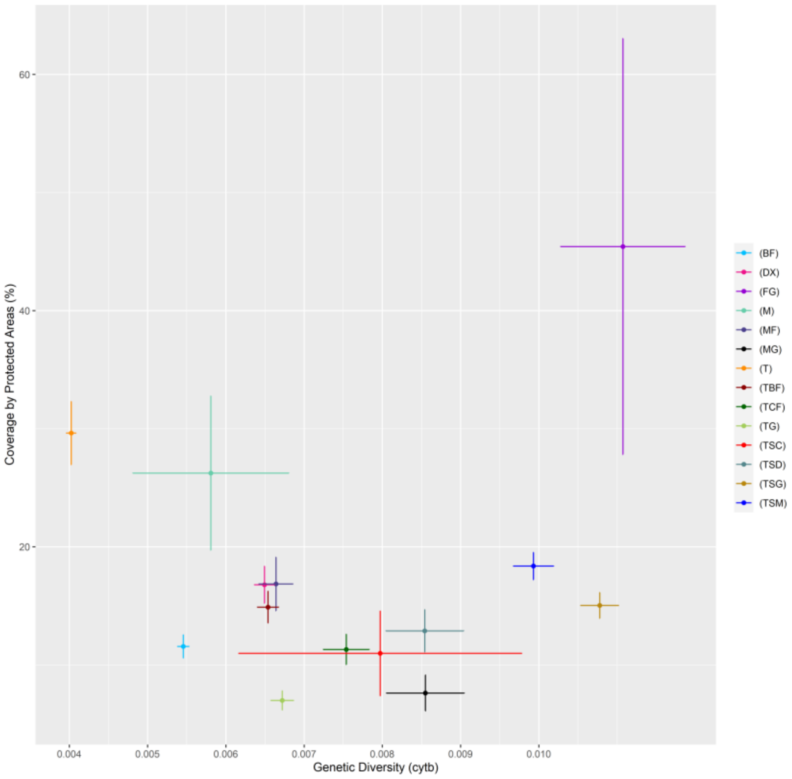


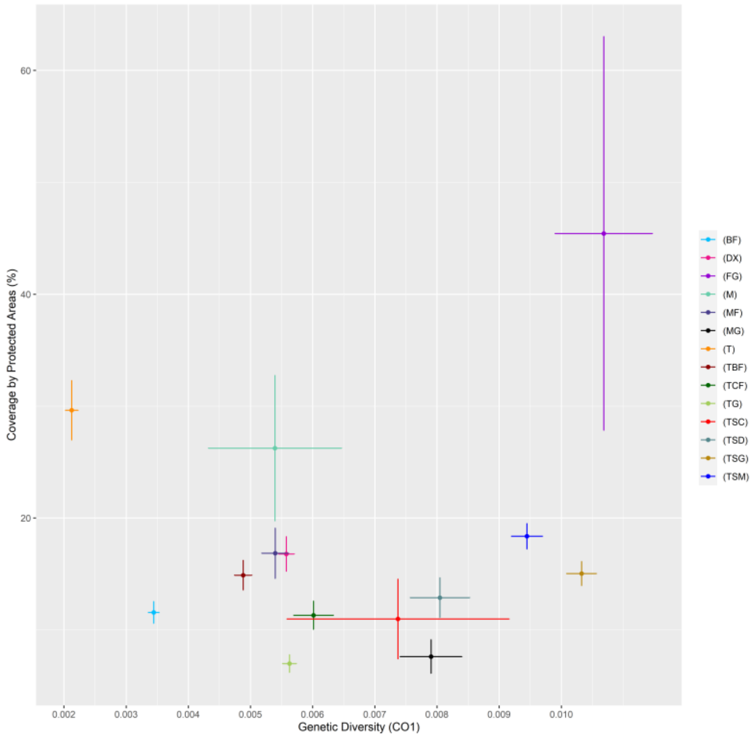


(b)

**Figure S3:** The mean coverage by protected areas as well as the mean of genetic diversity in each of the bioclimatic regions were calculated and were visualized using standard error.

**Figure S4: The geographical distribution of the residuals between genetic diversity (CO1) and species richness**

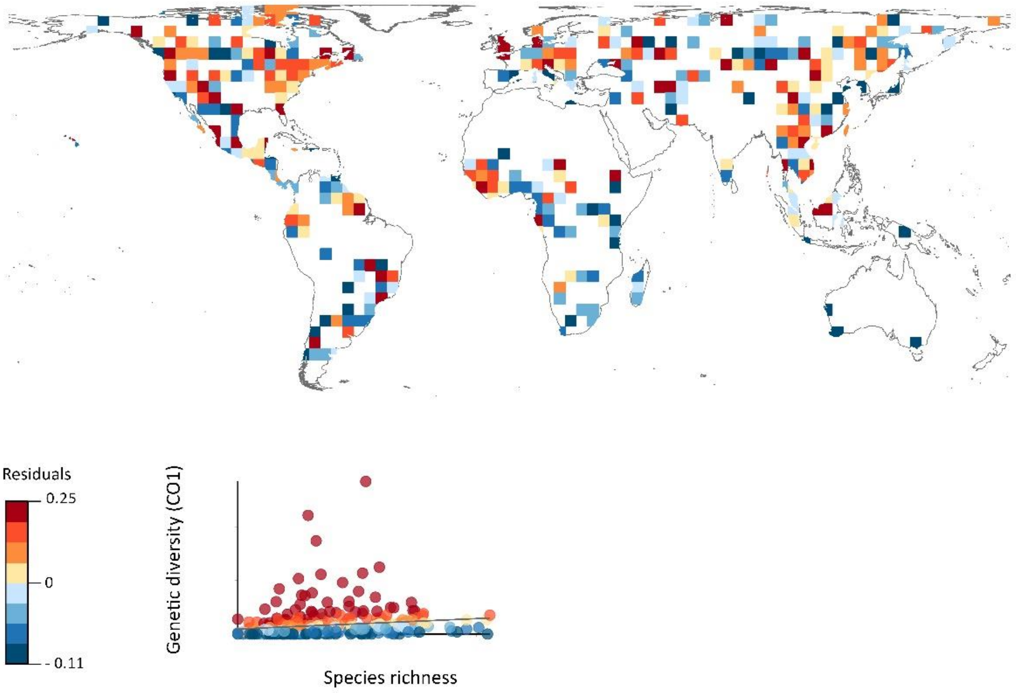


**Figure S4:** The map shows the geographical distribution of the residuals between genetic diversity (CO1) and species richness. Dark red visualized areas where genetic diversity is higher than predicted based on the species richness. Dark blue regions indicate the opposite, genetic diversity is lower than predicted through species richness.

**Table S1:** The relation between the coverage of protected areas and genetic diversity was tested for each bioclimatic region using simple linear regression analysis, whereby mangroves, flooded grassland and tropical, and subtropical coniferous forests were excluded due to low sample sizes.

(a) cytb

| *Region* | *Sample size* | *P value* | *Multiple R^2* | *Adjusted R^2* | *Multiple r* | *Adjusted r* | *95% CI* |
|---|---|---|---|---|---|---|---|
| TBF | 142 | 9.88e-10 | 0.2349 | 0.2294 | 0.484665 | 0.478957 | -966.99, -518.87 |
| T | 148 | 0.00104 | 0.07131 | 0.06494 | 0.267039 | 0.254833 | -2402.95, -619.08 |
| BF | 142 | 0.0469 | 0.02792 | 0.02098 | 0.167093 | 0.144845 | -615.5, -4.36 |
| MF | 50 | 0.3048 | 0.02193 | 0.001549 | 0.148088 | 0.039357 | -711.93, 227.36 |
| DX | 224 | 0.00108 | 0.0471 | 0.04281 | 0.217025 | 0.206906 | -690.11, -175.28 |
| TG | 84 | 0.2222 | 0.01812 | 0.006145 | 0.134611 | 0.07839 | -293.01, 69.1 |
| TCF | 28 | 0.517 | 0.01635 | -0.02148 | 0.127867 | | -409.6, 211.07 |
| TSM | 292 | 1.28e-07 | 0.09187 | 0.08874 | 0.303101 | 0.297893 | 158.28, 338.96 |
| TSG | 174 | 1.14e-07 | 0.1512 | 0.1463 | 0.388844 | 0.382492 | 232.28, 489.7 |
| MG | 32 | 0.341 | 0.03032 | -0.002007 | 0.174126 | | -108.32, 303.71 |
| TSD | 55 | 0.729 | 0.002278 | -0.01655 | 0.047728 | | -141.87, 201.41 |

(b) CO1

| *Region* | *Sample size* | *P value* | *Multiple r^2* | *Adjusted r^2* | *Multiple r* | *Adjusted r* | *95% CI* |
|---|---|---|---|---|---|---|---|
| TBF | 142 | 4.84e-06 | 0.1391 | 0.133 | 0.372961 | 0.364692 | -651.85, -269.87 |
| T | 148 | 0.986188 | 2.06e-06 | -0.006847 | 0.001435 | | -354.32, 348.16 |
| BF | 142 | 0.566 | 0.002362 | -0.004764 | 0.0486 | | -134.4, 244.84 |
| MF | 50 | 0.00182 | 0.185 | 0.168 | 0.430116 | 0.409878 | -1040.95, -252.82 |
| DX | 224 | 0.030097 | 0.02101 | 0.0166 | 0.144948 | 0.128841 | -523.31, -26.72 |
| TG | 84 | 0.2444 | 0.01649 | 0.004494 | 0.128413 | 0.067037 | -355.42, 91.82 |
| TCF | 28 | 0.4940 | 0.01817 | -0.01959 | 0.134796 | | -343.52, 170.17 |
| TSM | 292 | 3.81e-07 | 0.08522 | 0.08206 | 0.291925 | 0.286461 | 147.03, 326.25 |
| TSG | 174 | 2.8e-07 | 0.1426 | 0.1376 | 0.377624 | 0.370945 | 219.71, 476.76 |
| MG | 32 | 0.338 | 0.03059 | -0.001724 | 0.1749 | | -104.44, 294.51 |
| TSD | 55 | 0.845 | 0.000723 | -0.01813 | 0.026892 | | -155.26, 188,87 |

**Figure S5: The exposure to agriculture expansion in regions with low protection coverage (< 10 %) but high genetic diversity (belonging to the 20 % most diverse areas) under the four main SSP scenarios**

(a)

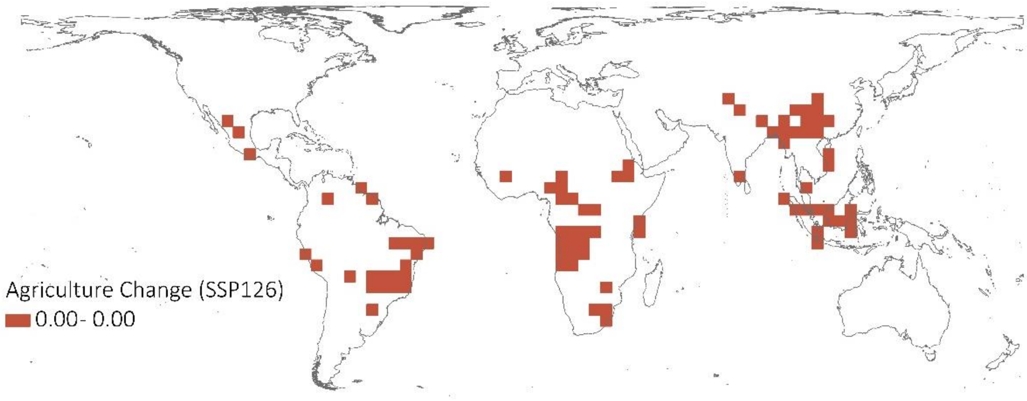


(b)

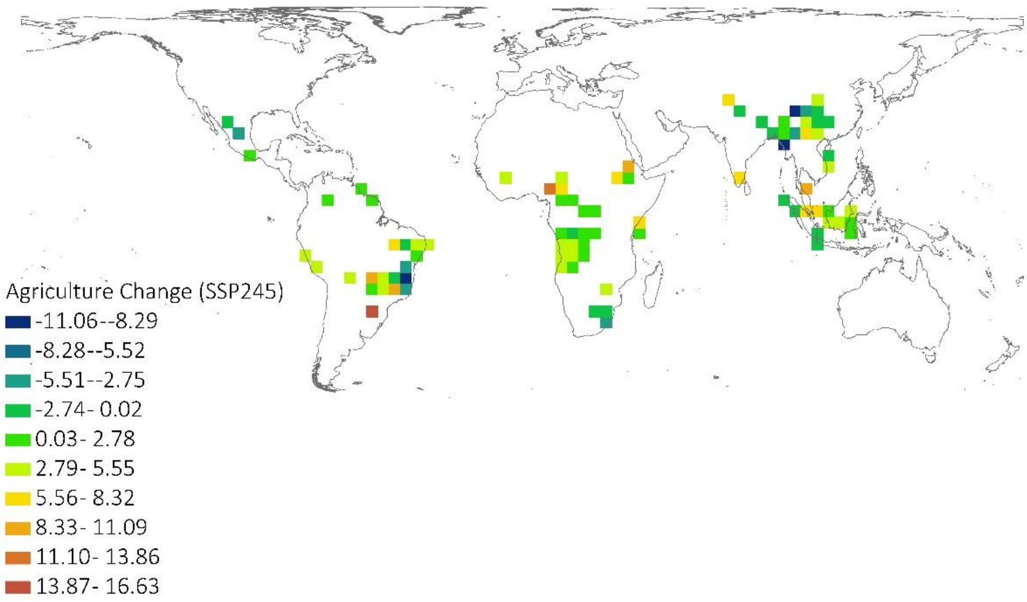

(c )

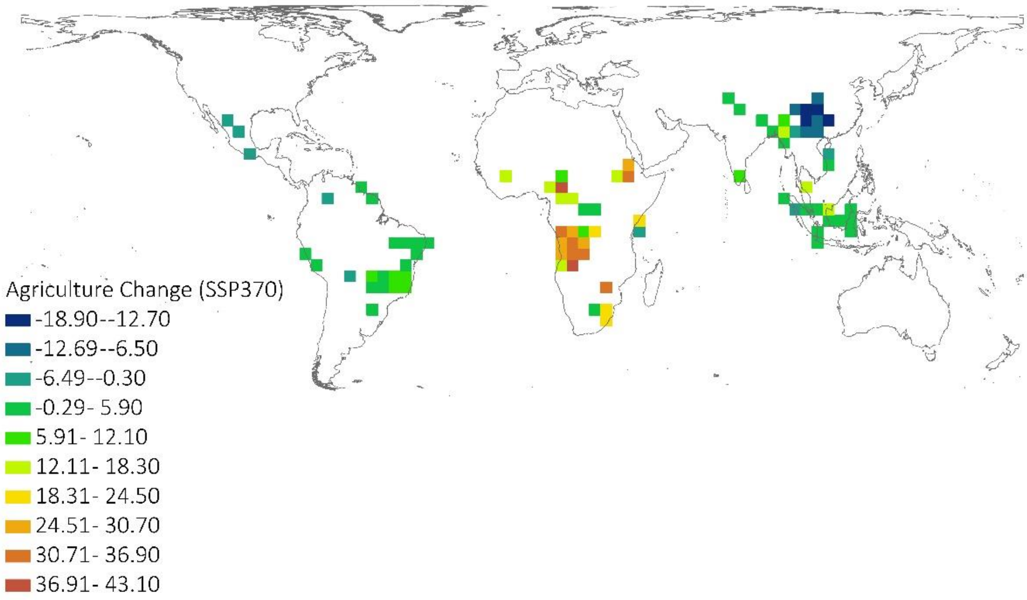
Agriculture Change (SSP370)
-18.90--12.70
-12.69--6.50
-6.49--0.30
-0.29- 5.90
5.91- 12.10
12.11- 18.30
18.31- 24.50
24.51- 30.70
30.71- 36.90
36.91- 43.10

(d)

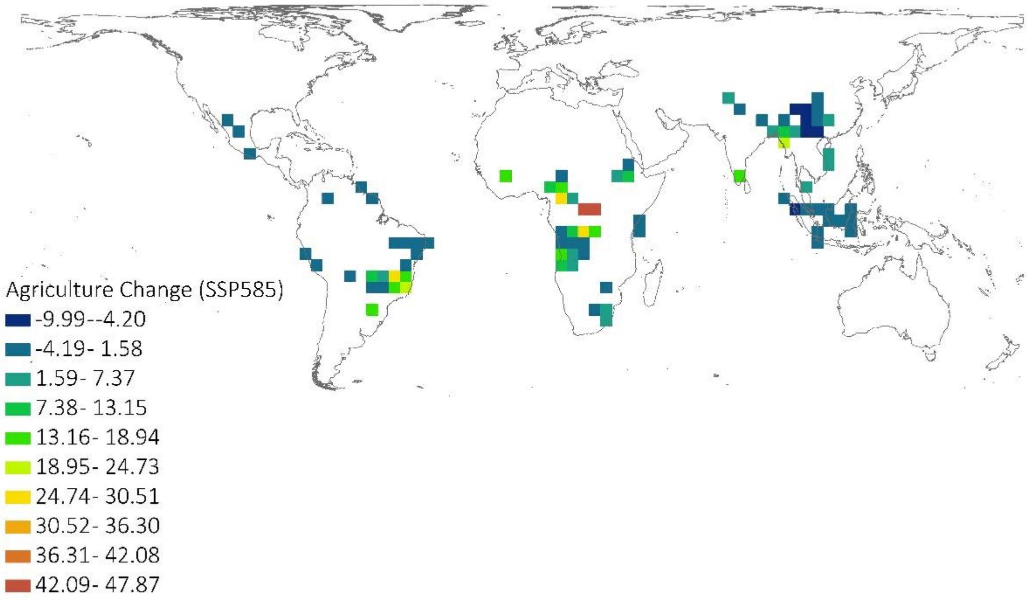
Agriculture Change (SSP585)
-9.99--4.20
-4.19- 1.58
1.59- 7.37
7.38- 13.15
13.16- 18.94
18.95- 24.73
24.74- 30.51
30.52- 36.30
36.31- 42.08
42.09- 47.87
